# The lateral intraparietal area preferentially supports stimulus selection in directed tasks compared to undirected free behavior

**DOI:** 10.1101/2022.03.09.483625

**Authors:** W. Jeffrey Johnston, Stephanie M. Tetrick, David J. Freedman

## Abstract

Our reactions to the sensory world depend on context. For instance, explicit directions to search for a particular object or feature will induce a different treatment of the sensory world than exploration without a fixed goal. To understand how we navigate the sensory world, it is necessary to understand how both directed search and undirected exploration are produced by the brain. The lateral intraparietal area (LIP) in the posterior parietal cortex has an established role in visual stimulus selection. However, most studies of LIP’s role in stimulus selection focus primarily on highly trained, directed tasks, in which animals are given explicit cues as to which stimulus they should select. Here, we compare neural activity in LIP across two tasks. In one task, the animal is given an explicit direction to select one of two natural images from an array; in the other, the animal is allowed to choose an image freely based on their innate preferences. We find that LIP reliably encodes the eye movement prior to its execution only in the directed task, while the eye movement encoding in the undirected task emerges significantly later. Further, LIP’s encoding of the image’s behavioral relevance emerges after the decision in both tasks. These results indicate that LIP preferentially supports stimulus selection in highly trained and directed behaviors, as opposed to free behavior.

## 1 Introduction

The same sensory input can have vastly different semantic meaning and urgency depending on the context in which it is received. For instance, a hiker might not ordinarily notice or react to small trickles of water seeping out of nearby rock; however, if the hiker is out of water, those same seeps of water become both extremely salient and evoke a strong behavioral response – as they might indicate a spring of fresh, potable water. This kind of search typifies directed stimulus selection. The hiker has a particular need, and they are searching their environment for ways to satisfy that need. On the other hand, without a specific need, the hiker will spontaneously explore the visual scene, attending to different parts of the scene according to a constellation of different factors – including low-level visual salience (e.g., contrast in color or spatial frequency)[1, 2] and remembered associations (e.g., familiarity)[3–6]. We refer to this kind of spontaneous free viewing as undirected stimulus selection.

Extensive previous work has studied the neural mechanisms underlying directed stimulus selection in non-human primates – much of it using explicit search tasks[7–10]. These studies have shown neural correlates of the search process in both prefrontal cortex (PFC; especially the frontal eye fields, FEF)[1, 10, 11] as well as in the lateral intraparietal area (LIP) of the posterior parietal cortex (PPC)[10, 12, 13]. In these regions, there is thought to be both a representation of the integrated behavioral priority – including task-relevance, low-level salience, and familiarity[12–15] – of different parts of visual space[16] as well as neural activity corresponding to the motor action itself[17]. Within this literature, increasing the low-level salience of the target (distractor) stimulus has been shown to make the task easier (harder) for the animal to perform[10]. Further, it has been shown that the LIP appears to play a preferential role when the animal’s decision is directly related to low-level salience, while the FEF and other prefrontal regions appear to play a preferential role when low-level salience is matched across stimuli and the animal uses a serial search procedure[10, 18, 19]. Due to this putative role in visual stimulus selection based on low-level salience, we hypothesize that LIP may also be involved in undirected stimulus selection, where stimulus selection is uncoupled from both explicit task demands and reward.

However, parallel work has shown that representations in LIP are strongly shaped by training[20–22] and behavioral context[23]. As a result of training on a complex task, LIP develops representations of abstract, task-relevant factors[20, 22], and is shown to have a causal role in task performance[24]. Further, these learned stimulus representations were stronger and more closely coupled to behavior than similar representations observed in dorsolateral prefrontal cortex[25]. These results show that extensive training can shape the role that LIP plays in complex behaviors, particularly by increasing its representation of different stimulus features. Thus, it is not clear that LIP’s established role in representing the behaviorally relevant stimulus features and mediating stimulus selection in highly trained and directed tasks generalizes to a role in stimulus selection in undirected, free-viewing contexts.

Here, we compare the activity of the same population of LIP neurons across two distinct tasks, which are performed in randomly interleaved blocks during the same sessions. In both tasks, monkeys make a saccade to one of two image stimuli. However, the tasks vary in the factors that drive the animals’ choice. In particular, one task is highly trained and directed, the saccade delayed match-to-sample task (sDMST) – mirroring the directed tasks widely used in the literature. The second task, however, is untrained and undirected, the preferential looking task (PLT). While the animal consistently views the images in the PLT, they are not rewarded based on their viewing behavior. Thus, these two tasks allow us to compare the role of LIP in stimulus selection with identical stimuli and very similar motor actions, but vastly differing behavioral and cognitive contexts. Because LIP has an established role in oculomotor actions per se[12, 26–28] and a putative role in stimulus selection based on both low-level and task-relevant[13, 29] stimulus features, we hypothesized that LIP would be strongly involved in both tasks. However, this is not what we find. First, we find stronger and earlier representations of both the saccade direction and the behavioral relevance of the images during performance of the sDMST relative to the PLT. In particular, a reliable representation of the saccade direction emerges prior to the first saccade in the sDMST, but not in the PLT. Second, in both tasks, we find that the representation of the behavioral relevance of the images emerges after the animal’s choice has been made. Thus, while LIP may be involved in saccade production during the sDMST, we do not find evidence that it provides a representation of behavioral priority that would be useful for decision-making in either task.

## 2 Results

### 2.1 Undirected and directed tasks with the same stimuli and behavioral response

To probe differences in engagement of LIP during undirected and directed stimulus selection, we trained two Rhesus macaques to perform two tasks in which the animals viewed natural image stimuli. The first task, the PLT, allows the animal to choose which image to view without external, trained constraints – and the animal was rewarded so long as they correctly fixated at the beginning of the trial (for more details, see *Behavioral tasks and training* in *Methods*; Figure 1b). Importantly, despite the absence of a reward contingency in the PLT, both animals engaged with the images. In particular, both animals used their first saccade to look at one of the two images on almost every trial (Monkey R: 98%, Monkey N: 96%) and they continued to view the two images for most of the remaining free viewing period (Monkey R: fixated on one of the two images for 84% of the free viewing time; Monkey N: 94%).

**Figure 1:**
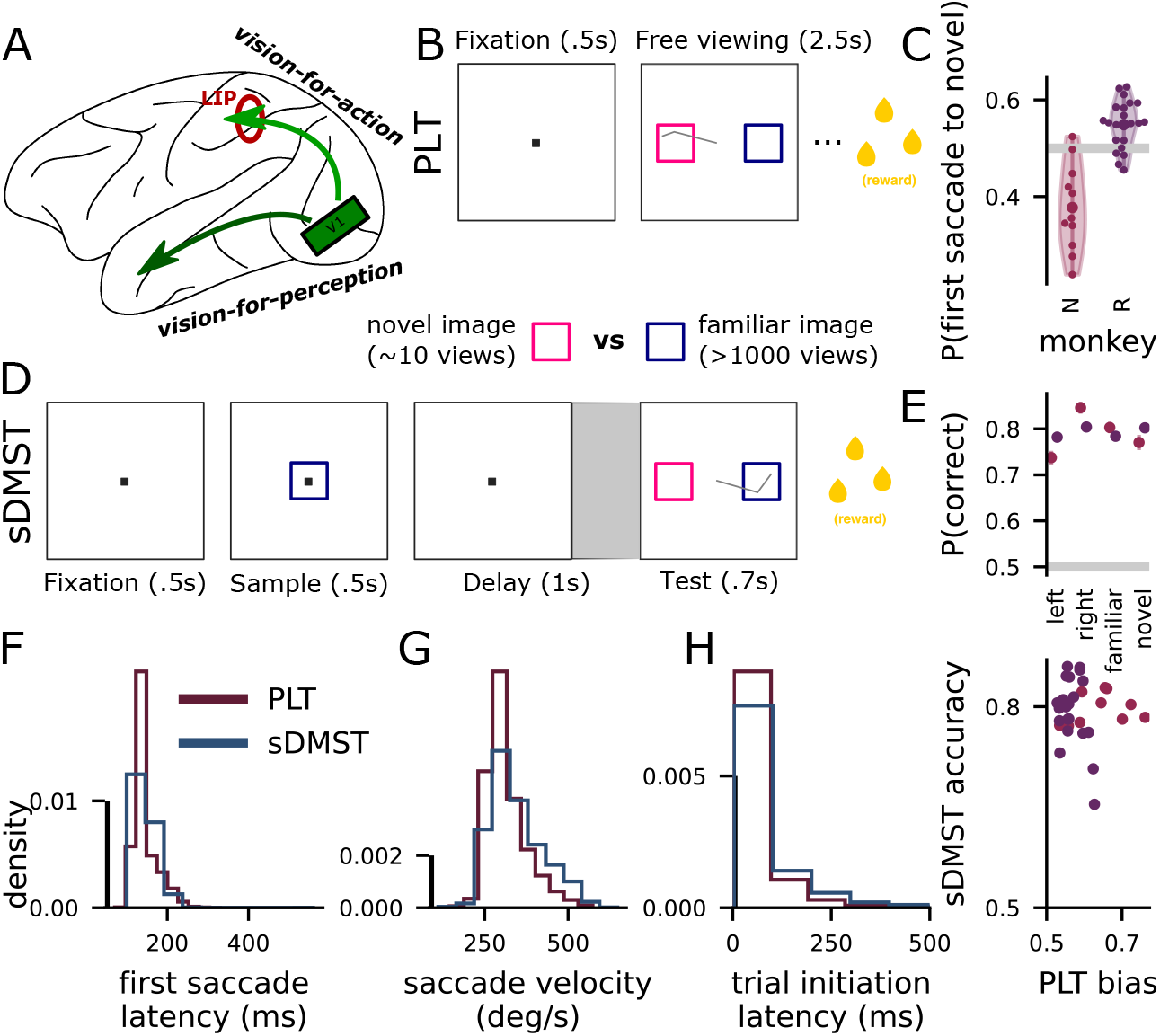
Task schematic and behavioral quantification. **A** Schematic of the Macaque visual system. Recordings were performed in the dorsal stream region LIP. **B** Schematic of the undirected preferential looking task (PLT). (below) The same novel and familiar images are presented in both tasks. **C** Violin plots of the distribution of first saccade biases in the PLT across experimental sessions. Both animals have reliable biases; Monkey R (purple) prefers novel images; Monkey N prefers familiar images. **D** Schematic of the directed saccade delayed match-to-sample task (sDMST). The images are presented in the same locations in both tasks. **E** Behavioral performance across different trial conditions. **F** Histogram of first saccade latencies. **G** Histogram of first saccade velocities. **H** Histogram of trial initiation latencies.

To influence the animal’s behavior on this undirected task, half of the natural images shown are from a set of novel images that is new to the animal at the start of each experimental session and the other half of the images are from a set of familiar images that the animal has seen over one thousand times previously (see *Image sets* in *Methods*). This manipulation of image familiarity affected the behavioral priority of the images for both animals, as demonstrated in their behavior. The two animals had different preferences: for the first animal, their first saccade was significantly more likely to go to a novel rather than familiar image (95% confidence interval for Monkey R: 52 - 56%, bootstrap test: *p* < .001); for the second animal, their first saccade was significantly more likely to go to a familiar rather than novel image (for Monkey N: 59 - 64%, bootstrap test: *p* < .001; see Figure 1c for these biases on individual sessions). Further, while this first saccade bias is modest, across all PLT trials both animals spent on average over 30% more time viewing images from their preferred set (novel or familiar) in the first 100 - 350 ms of the free viewing period than they spent viewing images from their nonpreferred set (Monkey R: 26 - 44% more time on novel images across sessions, bootstrap test: *p* < .001; Monkey N: 31-64% more time on familiar images across sessions, bootstrap test: *p* < .001; see Figure S.1b). This large difference in viewing time, coupled with the moderate difference in first saccade choice, indicates that there was a clear difference in the behavioral priority of the two kinds of images – and that this difference in priority exists prior to the first saccade. As an aside, to emphasize effect sizes in addition to statistical significance, we report bootstrapped 95% confidence intervals in conjunction with p-values throughout the paper.

The second task, the sDMST, explicitly constrains the animal’s choice through extensive task training. In particular, the animal must choose to look at the sample image they saw earlier in the trial to receive a reward (for more details, see *Behavioral tasks and training* in *Methods*; Figure 1d). After training and during recordings, both animals performed this task with 79 - 80% mean accuracy (Figure 1e). Importantly, while the temporal sequence of events differs across both tasks, at the start of the test period of the sDMST, the task is visually identical to the PLT at the start of the free viewing period, and the animal performs the same physical action as in the PLT: They make an eye movement to one of the two presented images. Across the two tasks, the animals’ choices are influenced by what we refer to as the behavioral priority of the images. In the sDMST, priority is related to the match-status of an image (i.e., match images have higher behavioral priority than non-match images); in the PLT, as mentioned above, priority is related to the familiarity of the images (and potentially incorporates other factors such as the salience of image features, see *The effect of low-level salience on behavior* in *Supplement*).

Despite the differences in task demands, our analyses of the animals’ behavior indicate that both animals were engaged in both tasks. In particular, while the latency of the animal’s first saccade in the free viewing period in the PLT and the test period was significantly later in the PLT relative to the sDMST in both monkeys (Monkey R: 134 ms in the PLT, 132 ms in the sDMST, 1 - 3 ms later in the PLT, bootstrap test: *p* < .001; Monkey N: 197 ms in the PLT, 170 ms in the sDMST, 24 - 29 ms later in the PLT, bootstrap test: *p* < .001; Figure 1f), both differences were small. Further, the mean velocity of the first saccade was slightly higher in the sDMST than the PLT for Monkey R (PLT: 314 ° s^−1^; sDMST: 358 ° s^−1^; 41 - 48 ° s^−1^ faster in the sDMST, bootstrap test: *p* < .001), while Monkey N had no significant difference (PLT: 321 ° s^−1^; sDMST: 321 ° s^−1^; −4.58 - 5.85 ° s^−1^ slower in the sDMST, bootstrap test: *p* = .381; Figure 1g). Finally, we investigated the latency to trial initiation as another proxy of behavioral engagement. Monkey R showed no difference in initiation latency (PLT: 35 ms; sDMST: 32 ms; difference: –1 - 6 ms, bootstrap test: *p* = .094; note that trials in which the animal is already fixating in the fixation window at the beginning of the trial are counted as zero latency to initiation) and Monkey N showed only a small difference (PLT: 107ms; sDMST: 139ms; 11 - 50 ms slower for the sDMST, bootstrap test: *p* < .001; Figure 1h). Thus, while there are small differences across the two tasks in these behavioral metrics, the animals were engaged in both tasks.

### 2.2 Single neurons are more active in the directed matching task

While the animals performed both the sDMST and PLT in randomized blocks of 20 trials per task, we recorded from 343 LIP neurons (Monkey R: 156, Monkey N: 187). These neurons were recorded across 22 and 11 recording sessions in Monkey R and N, respectively. In both monkeys, the majority of neurons had visual or saccade-related responses (that is, a significant firing rate change between the fixation period and stimulus presentation periods for at least one task, bootstrap test, *p* < .05; Monkey R: 153 neurons, Monkey N: 125 neurons; 143 and 84 neurons were selective in the PLT for Monkey R and N, respectively; 147 and 104 in the sDMST).

First, we investigated the responses of single neurons around the time of the first saccade (the saccade indicating the animal’s decision in the sDMST and the first saccade to an image in the PLT). Neurons show a variety of dynamics, with some showing elevated activity prior to the saccade (Figure 2a, b, bottom and top, respectively) and others peaking strongly only after the saccade (Figure 2a, b, top and bottom, respectively). Additionally, some neurons show higher firing in the PLT (Figure 2a, b, top and bottom) or sDMST (Figure 2a, b bottom and top; during the fixation period, Monkey R: 41/152 neurons fire significantly more in the sDMST and 34/152 fire significantly more in the PLT, Monkey N: 29/149 and 19/149; in the 200 ms centered on the first saccade, Monkey R: 56/152 neurons fire significantly more in the sDMST, 18/152 fire significantly more in the PLT, Monkey N: 45/149 and 16/149). Averaging across the population, we find a trend toward higher firing rates during the fixation period (Monkey R: 0 - 0.14z-scored higher in the sDMST, bootstrap test: *p* = .03; Monkey N: −0.04 - 0.06z-scored, bootstrap test: *p* = .294; Figure 2c, left) and significantly increased firing rates during the peri-saccade interval (Monkey R: 0.09 - 0.23z-scored higher in the sDMST, bootstrap test: *p* < .001; Monkey N: 0.05 - 0.18z-scored, bootstrap test: *p* < .001; Figure 2c, right). Thus, while there may be weakly elevated firing rates in the sDMST relative to the PLT during the pre-saccade fixation, there is a more reliable difference in firing rates during the saccade itself.

**Figure 2:**
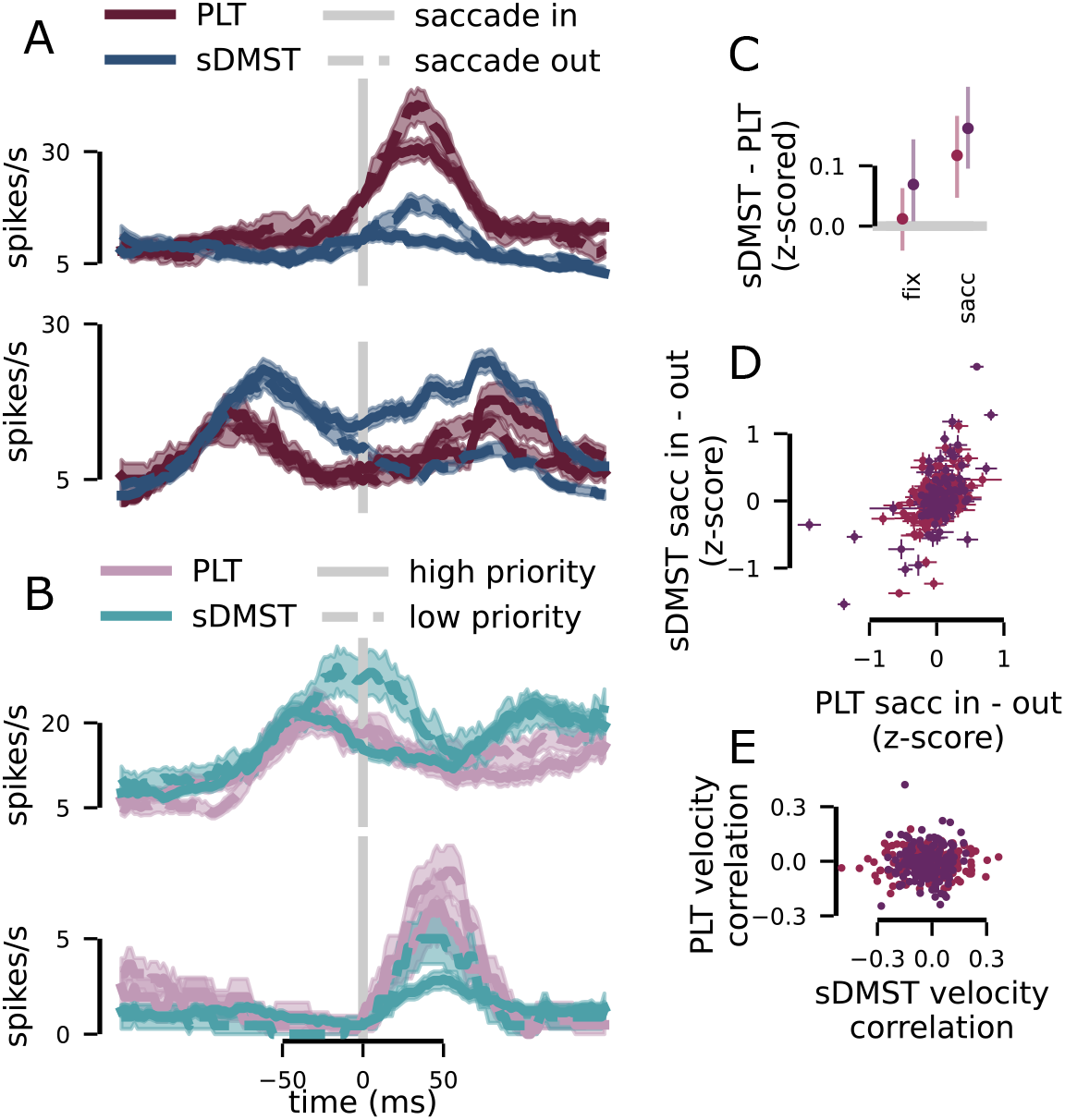
Single neurons are responsive in both the sDMST and PLT. **A** The activity of two example neurons, one from Monkey R (top) and the other from Monkey N (bottom). The traces contrast saccades into and out of the recorded hemifield across the two tasks. Time is centered around the animal’s first saccade in both tasks. **B** As in **A**, but contrasting when novel/familiar and match/non-match images are presented in the recorded hemifield. **C** The average z-scored firing rate difference between the sDMST and the PLT in the fixation period just prior to the first saccade (left, –200 - 0 ms before the fixation point turns off) and around the time of the animal’s first saccade (right, –100 - 100ms after the first saccade). **D** The average z-scored firing rate difference between saccades into and out of the recorded hemifield for each neuron. **E** The correlation between z-scored firing rate and saccadic eye movement velocity for each neuron, for both the PLT and sDMST.

Next, we asked whether there was a difference in firing rates based on the direction of the saccade – that is, whether the saccade was into the contralateral or ipsilateral hemifield (with respect to our recordings). Averaging across the population during the peri-saccade period, there was slightly – but not significantly – higher firing rates for saccades into the contralateral hemifield relative to saccades into the ipsilateral hemifield in both the sDMST (Monkey R: −0.04 - 0.14z-scored more firing for saccades into the contralateral hemifield, bootstrap test: *p* = .141; Monkey N: −0.02 - 0.08z-scored, bootstrap test: *p* = .138) and the PLT (Monkey R: −0.06 - 0.09z-scored more firing for saccades into the contralateral hemifield, bootstrap test: *p* = .3; Monkey N: −0.03 - 0.05 z-scored, bootstrap test: *p* = .274). Across the population of neurons, however, the difference in firing for the two saccade choices in the sDMST was correlated with the difference in the PLT (Monkey R: *r* =0.38 - 0.7, bootstrap test: *p* < .001; Monkey R: *r* =0.32 - 0.56, bootstrap test: *p* < .001; Figure 2d). Thus, the individual neurons in the population have consistent saccade tuning across the two tasks, but the population itself does not show a strong average difference in firing between the saccade directions.

Finally, we tested whether the moderately higher saccade velocities in the sDMST exhibited by Monkey R were correlated with higher firing rates in LIP, as this could be a purely physical difference in the saccades that might increase saccade tuning in the sDMST relative to the PLT. In particular, we tested whether there was a reliable correlation between saccade velocity and pre-saccadic firing in both tasks across our population. In Monkey R, this revealed a small but significant negative correlation between saccade velocity and peri-saccade firing (r =-0.04 - −0.01, bootstrap test: *p* < .001) and no significant effect for the PLT (r =–0.02 - 0.01, bootstrap test: *p* = .383). The direction of the effect for the sDMST is inconsistent with the moderately increased saccade velocities explaining greater saccade tuning during the sDMST. For Monkey N, there was no significant effect for either case (sDMST: *r* =–0.05 - 0.01, bootstrap test: *p* = .044; PLT: *r* =–0.02 - 0, bootstrap test: *p* = .075). We further tested whether this correlation was reliable across the two contexts by assessing whether a neuron’s firing rate-saccadic velocity correlation in the sDMST predicts the firing rate-saccadic velocity correlation in the PLT. We found no significant relationship between the correlations across these two contexts (Figure 2e).

### 2.3 Choice encoding emerges earlier in the directed matching task

Our results at the single neuron level show that LIP has somewhat higher firing rates in the sDMST relative to the PLT primarily around the time of the animal’s saccade. To understand what this difference in single neurons means at the population level, we apply neural population decoders to characterize the time-course and magnitude of information about the animal’s saccade. We trained support vector machine (SVM) classifiers to decode saccade direction (that is, whether the saccade was into the contra- or ipsilateral hemifield) from neural activity during each of the two tasks. For all decoding analyses, training and testing trials are balanced across conditions and the decoding performance reflects performance on held out trials from 10-fold cross-validation (more details of these analyses are discussed in *Support vector machine (SVM) analyses* in *Methods*). In all comparisons across the PLT and sDMST, the same neurons and the same number of trials (*N* = 30) are used.

This analysis revealed that significant saccade direction decoding emerges earlier in the sDMST compared to the PLT (14 - 28ms earlier, bootstrap test: *p* < .001; Figure 3a, left). Further, decoding performance is significantly greater than chance prior to the first saccade for the sDMST, while decoding performance rises more slowly and asymptotes later in the PLT (from –50 - 0 ms relative to the first saccade, sDMST: 58 - 84%, bootstrap test: *p* = .004; PLT: 46 - 68%, bootstrap test: *p* = .162; decoding performance in the sDMST is from –3-32% higher, bootstrap test: *p* = .053; 218 neurons; Figure 3a, right). That is, the same neural population encodes the saccade direction more strongly in the directed sDMST than in the undirected PLT. This pattern of results holds when the analysis is repeated on only the neurons with the strongest visual responses. This pattern is further replicated across individual recording sessions as well, where saccade decoding from individual sessions just prior to the first saccade revealed reliably higher performance in the sDMST relative to the PLT (Monkey R: 52 - 59% in the sDMST, 46 - 53% in the PLT, 2 - 9% higher, bootstrap test: *p* < .001; Monkey N: 52 - 57% in the sDMST, 49 - 55% in the PLT, –1 - 6% higher, bootstrap test: *p* = .098; Figure 3b).

**Figure 3:**
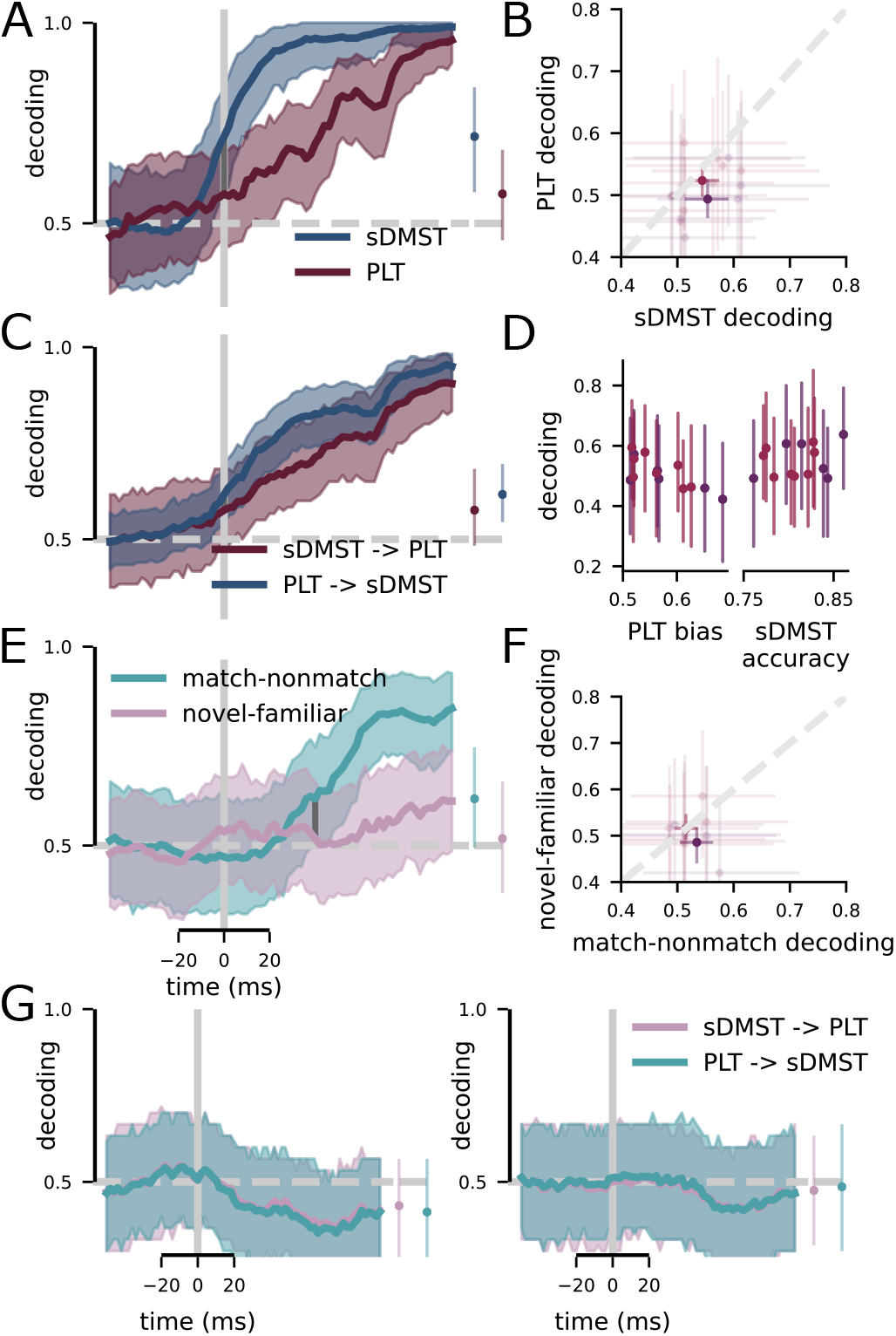
The same population of LIP neurons encodes more information about behavioral response and image relevance in the directed sDMST relative to the undirected PLT. **A** The decoding performance for SVM classifiers trained to determine whether the animal made a saccade into or out of the recorded hemifield, contrasted across both tasks. (left) Time-course of decoding; (right) decoding performance from –50 - 0 ms around the time of the first saccade. **B** Decoding performance in individual experimental sessions across both tasks prior to the first saccade (–50 - 0 ms, faded dots are individual sessions, solid dots are averages across sessions). **C** Cross-task decoding performance. In the red (blue) trace, classifiers were trained on data from the sDMST (PLT) and tested on data from the PLT (sDMST). Performance is similar and high going in both directions indicating a similar representation of saccades in both tasks. (left) Time-course of decoding; (right) decoding from –50 - 0 ms around the time of the first saccade. **D** (left) Saccade decoding performance during the PLT from a single experimental session compared to the magnitude of the animal’s familiarity bias on that session. (right) Saccade decoding performance during the sDMST from a single experimental session compared to the accuracy on the sDMST. **E** Decoding performance for SVM classifiers trained to discriminate between when the image in the recorded hemifield is either a match or nonmatch in the sDMST and novel or familiar in the PLT. (left) Time-course of decoding; (right) decoding performance from –10 - 40 ms around the time of the first saccade. **F** Decoding performance just after the first saccade in individual experimental sessions (– 10 - 40 ms, faded dots are individual sessions, solid dots are averages across sessions). **G** Cross-task decoding performance for priority for Monkey N (left) and Monkey R (right).

Next, we use a cross-decoding approach to ask whether neurons play similar roles in choice encoding across these two tasks by training the SVM classifier on trials from one task and then testing on trials from the other task. Consistent with LIP’s established role in saccade planning, cross-decoding from both the sDMST to the PLT and from the PLT to the sDMST produces performance that is similar to when the two classifiers are trained and tested on data from the same task (from – 50 - 0 ms, trained on sDMST and tested on PLT: 0.48 - 68%; trained on PLT and tested on sDMST: 55 - 70%; Figure 3c). This indicates that the representation of saccade direction is shared between tasks, and further suggests that the differences in saccade representations that we see across the two tasks are explained well by an increase in gain of saccade direction selectivity in the sDMST relative to the PLT.

Further, we asked whether the strength of saccade encoding across a particular experimental session would correlate with the animal’s behavioral performance on either the PLT or the sDMST in that session. In particular, we hypothesized that we may get stronger decoding results on sessions in which the animals either performed the sDMST with greater accuracy or had a stronger familiarity bias in the PLT. However, in both cases, we do not find evidence of a relationship between behavioral performance and per-session decoding performance (Figure 3d).

### 2.4 Priority encoding emerges earlier in the directed matching task

We again use SVM decoding to assess the strength and time-course of the neural representation of the behavioral priority of the images – that is, decoding match and non-match in the sDMST and familiarity and novelty in the PLT. In these analyses, the training and test sets have an equal number of trials from all combinations of behavioral priority and saccade choice, which removes any contribution to decoding from the previously characterized saccade direction encoding – see *Support vector machine (SVM) analyses* in *Methods* for additional details. This analysis reveals significant decoding for behavioral priority in the sDMST (i.e., decoding of match vs. non-match status; emerging 40 - 48 ms after the first saccade). In contrast, we do not detect significant decoding novelty or familiarity in the PLT. This is just after the animal’s decision but just before the animal receives feedback about the outcome of the trial in the sDMST or the second saccade in the PLT (from –10 - 40 ms after the first saccade, sDMST: 49 - 75%, PLT: 38 - 66%, difference: –9 - 29%, 147 neurons; Figure 3e). This pattern of results holds when the analysis is repeated on only the neurons with the strongest visual responses. This pattern of results weakly replicates across individual sessions as well (Monkey R: 50 - 56% in the sDMST, 44 - 52% in the PLT, –1 - 12% higher decoding in the sDMST, Monkey N: 49 - 54% in the sDMST, 49 - 55% in the PLT, –2 - 2% higher decoding in the sDMST; Figure 3f). These results are broadly consistent with the idea of LIP serving as a priority map[13] in directed tasks, where there is an explicit reward contingency on the animals’ choices, but less so in undirected tasks without explicit reward contingencies. However, in both cases, the priority map-like representation emerges after the animal has made their decision about which stimulus to look at. Thus, in these tasks, the map may primarily be useful for evaluation of decisions after they have been made, rather than decision making per se.

Next, we use cross-decoding analyses to quantify to what extent this representation of behavioral priority is shared or distinct across the two tasks. This cross-decoder does not reach statistical significance in either of our two animals, indicating that the shared representation of behavioral priority is weaker than the shared representation of saccade direction. In both monkeys, the cross-decoders trend toward significantly below-chance performance. This effect is strongest in Monkey N and consistent with Monkey N’s behavioral preference for familiar images. However, cross-decoding performance also trends toward slightly below chance in Monkey R, reflecting a potential mapping between familiar and match images in Monkey R as well, which is inconsistent with Monkey R’s behavioral preference for novel images.

### 2.5 Low-level stimulus features are also weakly encoded in the undirected task

Image familiarity relies on the animal’s past experience with the image, and is a factor that can impact the salience or priority of a stimulus both in general and specifically in the PLT. Signatures of image familiarity have been observed in several brain regions[30, 31] (including LIP[15]), and particularly in the inferotemporal cortex[32–34] and hippocampus[4]. Although LIP activity did not obviously reflect image priority due to familiarity in the PLT, it is possible that LIP may still act as a priority map, but only for low-level stimulus features, such as local contrast. To test this hypothesis, we included a third task condition in the experiments described above that was identical to the PLT except that the novel and familiar natural image sets were replaced with high- and low-contrast isoluminant squares of the same size as the images which were displayed in the same positions as in both other tasks(Figure 4a). We refer to this modified PLT as the contrast-based PLT (conPLT). In this task, only one of our two animals had a reliable contrast preference (Monkey R: 46 - 53% of first saccades went to the high contrast square, bootstrap test: *p* = .624; Monkey N: 51 - 60% of first saccades went to the low contrast square, bootstrap test: *p* = .022; Figure 4b). In addition, the latency of first saccades in the conPLT were significantly longer than in the sDMST or standard PLT (Monkey R: 179 ms in the conPLT, 132 ms in the sDMST, 41 - 54 ms longer in the conPLT, bootstrap test: *p* < .001; Monkey N: 254ms in the conPLT, 170 ms in the sDMST,76 - 94ms longer in the conPLT, bootstrap test: *p* < .001; Figure 4c). Thus, the animal may be significantly less engaged during the conPLT than the PLT or sDMST. However, both animals made their first saccade to one of the two squares on the majority of trials (Monkey R: 78%; Monkey N: 93%).

**Figure 4:**
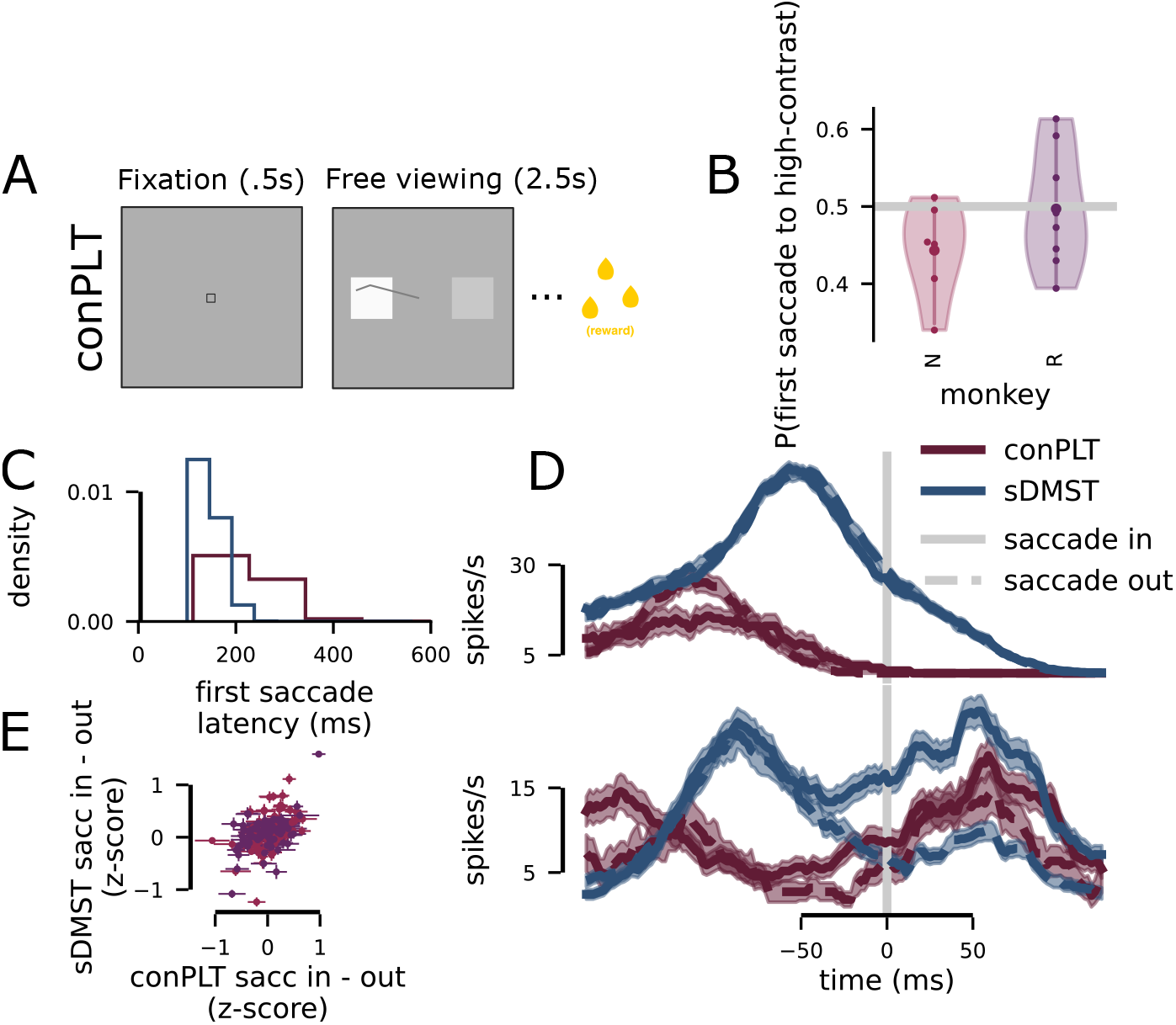
An undirected task where the visual stimuli are high- and low-contrast squares. **A** A variant of the PLT, which is referred to as the contrast preferential looking task (conPLT). It is the same as the PLT except that isoluminant squares with either high- or low-contrast are presented instead of natural images. **B** On different sessions, the animals have different biases for viewing either low (Monkey N, left) or high (Monkey R, right) contrast squares. **C** The latency of the animal’s first saccade is slightly greater in the conPLT compared to the sDMST. **D** Example neurons from Monkey R (top) and Monkey N (bottom); time is organized around the animal’s first saccade. **E** The average z-scored firing rate difference between saccades into and out of the recorded hemifield for each neuron.

Single neuron activity around the time of the first saccade during this task is similar to both of the other tasks (Figure 4d). Population average firing was significantly larger in the sDMST relative to the conPLT in Monkey R (0.24 - 0.47 z-scored, bootstrap test: *p* < .001) but only trended toward significance in Monkey N (–0.03 - 0.14 z-scored, bootstrap test: *p* = .093). There was no significant difference in either monkey during the fixation period. The saccade tuning of individual neurons, however, was consistent across the conPLT and sDMST (Monkey R: *r* =0.19 - 0.68, bootstrap test: *p* < .001; Monkey N: *r* =0.32 - 0.58, bootstrap test: *p* < .001; Figure 4e).

Next, we performed decoding analyses on these data to compare population encoding of choice as well as of match and contrast. Our population analyses reveal broad similarities between the relationship between the sDMST and the conPLT to those we have already established between the sDMST and the standard PLT. In particular, an encoding of the saccade direction emerges significantly later in the conPLT than the sDMST (conPLT: 80 - 82 ms after the first saccade, sDMST: 4 - 10 ms; 72 - 78 ms later in the conPLT, bootstrap test: *p* < .001; Figure 5a, left, 121 neurons, 20 trials per condition). On individual sessions, decoding performance tended to be higher in the sDMST than in the conPLT just after the first saccade (from −40 - 10 ms, combined: 0 - 5% higher in the sDMST than in the conPLT, bootstrap test: *p* = .029; Figure 5b). For encoding of match status and contrast, the results are less clear across the combined population (100 neurons, Figure 5c, d). However, trends are similar to those in the comparison between the sDMST and PLT (Figure 3).

**Figure 5:**
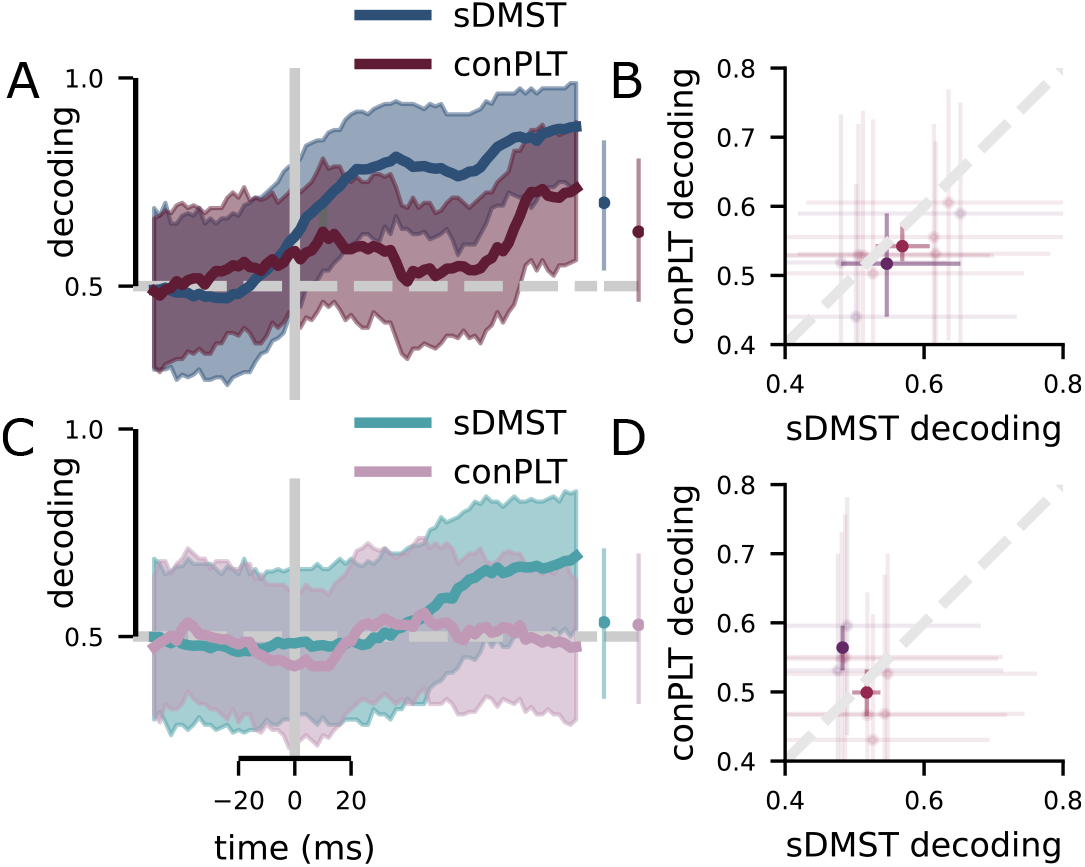
LIP is also less engaged in undirected behaviors that depend on low-level stimulus features. **A** The decoding performance of SVM classifiers trained to determine whether the animal made a saccade into or out of the receptive field (as in Figure 3a). (left) The time-course of decoding; (right) decoding performance −40 - 10 ms around the time of the first saccade. The pattern is similar to that shown for the PLT and sDMST. **B** Decoding performance for saccades in individual experimental sessions (−40 - 10 ms, faded dots are individual sessions, solid dots are averages across sessions); similar to Figure 3b. **C** The decoding performance of SVM classifiers trained to determine whether there was a match or nonmatch and high- or low-contrast square in the receptive field; similar to Figure 3a. (left) The time-course of decoding; (right) decoding performance 15 - 55 ms after the first saccade. **D** Decoding performance for image relevance in individual experimental sessions (15 - 55 ms, faded dots are individual sessions, solid dots are averages across sessions); similar to Figure 3f.

## 3 Discussion

In this study, we asked whether LIP’s established role in highly-trained and directed visual stimulus selection generalizes to untrained and undirected stimulus selection. In contrast to our expectation that LIP would play a generalized role in both these forms of stimulus selections, we found that LIP appears to be significantly less involved in stimulus selection during the undirected PLT than the directed sDMST, even when the physical saccades themselves, and the targets of those saccades, are similar across both tasks. LIP also appears less involved in the representation of image priority in the undirected than directed task. This pattern of results is replicated on an even simpler undirected preferential looking task where the animal chooses to view either a high- or low-contrast stimulus. Thus, the reduced task-relevant activity in LIP during the undirected task is likely not due to our choice of familiarity as the manipulated image feature in the PLT. While previous work has shown that the PPC responds more strongly during saccades made to visual stimuli than during saccades made in the absence of a visual target[35–37], our work shows that a similar difference exists between saccades during undirected and directed behavior – even when the same kind of visual stimulus is the target of both. This indicates that LIP may be more engaged in tasks that have explicit direction, while undirected tasks with similar overt behaviors may be mediated by other brain regions – perhaps those even more closely associated with oculomotor control, such as FEF or the superior colliculus.

We believe that these results illuminate one shortcoming of a common form of inference in neuroscience. In particular, many well-controlled experimental tasks are designed to mimic naturalistic behaviors. For instance, search tasks are often analogized to navigating in cluttered environments with many potential stimuli to select from. Then, neural signals are recorded in the highly trained task with explicit reward contingencies, and used to infer the roles of different brain regions in a core behavioral function, which is implicitly generalized to behavioral contexts without explicit reward contingencies. Our results point out a potential weakness of that inference step by showing that the same behavioral function – in our case, visual stimulus selection – has markedly different neural correlates in a trained, directed task relative to in an untrained, undirected task. These results emphasize the need to study spontaneous, “every day” behaviors (that do not require extensive training) alongside highly trained and well-controlled tasks to gain a more complete understanding of the neural correlates of core behaviors like stimulus selection.

Our results are surprising in the context of other studies on LIP using trained tasks. However, there were several other differences that may influence our findings. First, as noted earlier, experimental task training may increase the role LIP plays in a particular task, as reflected by LIP’s greater engagement in the highly trained sDMST relative to the untrained and naturalistic PLT. Second, in both of our tasks, the animal is allowed to make their behavioral response immediately upon presentation of the target images. This differs from many commonly used tasks, in which animals are trained to withhold their behavioral response until a “go” cue is provided. Thus, it is possible that LIP may be more involved in decisions where the animal chooses to deliberate due to task difficulty (e.g., [24]) or is forced to deliberate, rather than in the low-latency decisions made during both the sDMST and PLT (see Figure 1f). Finally, for most of our recordings (see *Electrophysiological recording* in *Methods* for more details), we used multi-site linear electrodes. While we used the standard memory-guided saccade task as a functional diagnostic for whether we were recording from LIP neurons (see *Memory-guided saccade* in *Methods*), we included neurons in our analysis that were in the same anatomical region, but that did not necessarily exhibit the characteristic memory saccade profile of an LIP neuron or have the same response field location[38]. As most of the existing literature has analyzed LIP neurons with this specific functional profile, we are likely analyzing a more diverse set of neurons than in many previous data sets in the literature.

Our results may also be surprising in the context of recent results showing that inactivation of PPC, including LIP, significantly affects how correlated an animal’s free viewing behavior is with the low-level salience of a natural image[19] (not due wholly to the hemifield neglect previously reported with LIP inactivation[9]). Those results indicate that LIP has a role in guiding the deployment of visual attention according to low-level image salience, outside of a directed task context. However, there are several key differences between the two studies. First, our main free viewing task did not rely on low-level visual salience, derived wholly from features like contrast and spatial frequency, as used in the other study – instead, we used an experience dependent and memory-related image familiarity signal to influence the animal’s behavior. Thus, it is possible that PPC is more involved in the representation of low-level image salience than image familiarity, which is more abstract. However, if this were the case, we would expect our high- and low-contrast control task to have revealed that role – though, since the animals were not strongly engaged by our contrast stimuli, it is possible that LIP plays a role in representation of low-level salience when that salience is more behaviorally relevant (as in trained tasks or other behavioral contexts). Second, in our study, two smaller images were presented with 180° angular separation in the periphery and then freely viewed; in contrast, in the inactivation study[19], one large image was centrally presented for free viewing. Thus, it is possible that PPC has a specialized role in representing the relative salience of different spatial locations in close proximity within a single visual scene, consistent with its previously described role in spatial processing[39, 40] and its described role in pop-out visual search tasks[10]. However, in our experiments, we also included trials on which the two stimuli were presented with only 45° of separation in visual angle in both the PLT and the conPLT. We did not observe a significant difference in either familiarity or contrast decoding between this condition and the 180° separation condition (Figure S.3). While this does not precisely replicate pop-out conditions, it indicates that closer spatial arrangement alone is not enough to engage LIP in undirected free viewing tasks to the same degree as it is engaged in directed search tasks.

There are two alternative ways to reconcile the conflicting conclusions from our study and the inactivation study discussed above[19]. First, the disruption to salience-related behavior may be due to a generalized disruption of visual – or, in particular, spatial[39, 40] – processing that results from the inactivation of PPC. Second, the effect on salience preference reported in the inactivation study[19] is relatively small, and indicates that LIP may not be the only, or even the primary, region involved in producing that salience preference. Thus, LIP may play a minor role in producing salience-related behavior that our experiment here did not detect. However, if this is true, our conclusion that LIP is more involved in representing image relevance in a directed rather than undirected context still holds. While our study provides correlational evidence that LIP plays a reduced role in undirected visual behaviors, further experiments could verify that role by inactivating LIP in directed and undirected tasks similar to the ones used here. We predict that inactivation will disrupt behavior to a larger extent in the directed rather than undirected task.

This work illustrates the importance of studying and comparing neural function in a variety of both naturalistic and highly-controlled behavioral tasks. Together with previous work, our results highlight the malleability of LIP as a result of extensive behavioral training and suggest that LIP may be preferentially involved in directed tasks with explicit and learned behavioral contingencies. This conceptualization of LIP makes an interesting prediction: That LIP will only mediate saccades toward and preferentially represent pop-out stimuli (as in [10]) when the animal has been trained to respond to those stimuli in an explicit task. While pop-out stimuli are still expected to capture attention without explicit task training, our results suggest that LIP would have a reduced role in mediating attention-related behaviors without explicit training.

## Acknowledgments

This work was supported by NIH F31EY029155 (WJJ), NSF 1707398 (WJJ), Gatsby Charitable Foundation GAT3708 (WJJ), NIH R01EY019041 (DJF), CRCNS NIH R01MH115555 (DJF), NSF NCS 1631571 (DJF), and a DOD Vannevar Bush Fellowship (DJF).

## Author contributions

WJJ and DJF conceived of the project and developed the experiments. WJJ and SMT trained the animals and performed the electrophysiological recordings. DJF supervised the experimental work. WJJ analyzed the data and made the figures. WJJ and DJF wrote the paper.

## 4 Methods

### 4.1 Surgical preparation and experimental setup

Two male monkeys (Macaca mulatta; Monkey R: 10 years old, ~10 kg; Monkey N: 14 years old, ~14 kg) were implanted with head posts and then trained on four behavioral tasks described below. Monkey R was implanted with a recording chamber positioned over PPC after training was complete. Monkey N had a recording chamber positioned over LIP from a previous series of motion categorization experiments that was used in this set of experiments as well. He had the chamber during training and it was maintained with regular sterile cleaning throughout. Our surgical, behavioral, and neurophysiological approach has been described in detail previously[1, 2]. Stereotaxic coordinates for chambers placement were determined using structural magnetic resonance imaging (MRI) that was performed prior to the headpost implantation to avoid artifacts. Both chambers were centered on the intraparietal sulcus. In Monkey R, the chamber was centered over the right cortical hemisphere, 23.8 mm lateral from the midline and 1 mm posterior to the intra-aural line. In Monkey N, the chamber was centered over the right cortical hemisphere, 9 mm lateral from the midline and 5.25 mm posterior to the intra-aural line. Both monkeys were housed in individual cages with a 12 hour light-dark cycle. Behavioral training and recordings always occurred during the light part of the cycle. During behavioral training and recordings, monkeys sat in custom-made primate chairs and were head-fixed. Task stimuli were displayed on a 21-inch color CRT monitor (1280*1024 resolution, 75 Hz refresh rate, 57 cm viewing distance). Identical timing and rewards were used for both monkeys. Different image sets were used for across the different monkeys and the details of this are described below, in *Image sets* in *Methods*. A solenoid-operated reward system was used to deliver juice reward to the monkeys. Monkeys’ eye positions were monitored by an optical eye tracker (SR Research) at a sampling rate of 1 kHz and stored for offline analysis. Stimulus presentation, task events, rewards, and behavioral data acquisition were accomplished using an Intel-based PC equipped with MonkeyLogic software running in MATLAB (http://www.monkeylogic.net)[3, 4]. All experimental and surgical procedures were in accordance with the University of Chicago Animal Care and Use Committee and National Institutes of Health guidelines.

### 4.2 Behavioral tasks and training

The four behavioral tasks used in this work are described here. The PLT and sDMST are our primary experimental tasks. The memory-guided saccade task was used to functionally probe for LIP-like responses[5, 6]. The dimming detection task was used to familiarize both animals with their familiar image sets.

#### 4.2.1 Preferential looking task (PLT)

The trial begins with a 500 ms fixation period. After the fixation period, the fixation point disappears and two images appear. Each image is drawn from one of two sets of natural images: the novel image set or the familiar image set (see *Image sets* in *Methods*). The animal is allowed to freely view both images. After 2.5 s of free viewing, the animal is provided with fluid reward. This reward is only contingent on completion of the initial fixation period, not on behavior during the free viewing period.

#### 4.2.2 Saccade delayed match-to-sample task (sDMST)

The sDMST begins with a fixation period, followed by the foveal presentation of a sample natural image (500 ms). Then, the image disappeared and the animal had to maintain fixation for a 1 s delay period. At the end of the delay, an array of two test images appeared, one in each hemifield, on opposite sides of the fixation target (that is, with 180° of angular separation). One of the test images is always the same as the sample image. After the test images appear, the animal must saccade to the test image matching the sample image within 400 ms and then hold that fixation for at least 400 ms. If the animal completes this successfully, then they are given a liquid reward. If the animal makes a saccade to the non-matching image, then the trial is aborted and the images disappear.

Both animals were trained to perform the sDMST over six months to one year. First, both animals were trained on the task using solid color squares (red and green). Then, a reduced set of natural images was introduced and both animals were further trained. Finally, both animals were trained on the task with the set of familiar images used in the experiments and different sets of novel images until they reached consistent levels of performance on those image sets as well. In both animals, near the end of their training on the sDMST, they were trained on the dimming detection task, which was used to familiarize the animals with their familiar image set.

#### 4.2.3 Memory-guided saccade

The memory-guided saccade task[7] is used to establish whether recordings are made from LIP, as neurons in LIP are believed to exhibit characteristic response properties in this task[5]. In the task, the animal initiates a trial with a fixation period. Then, a small luminance target is flashed for 300 ms at one of eight locations, equally spaced in the periphery. After a 1000 ms delay period in which the animal must hold fixation, the fixation point disappears and the animal must make an eye movement to the location of the target flash. There is no target present on the screen at the time of the saccade. If the animal completes the saccade within 500 ms and holds fixation for 150 ms, then they receive reward.

LIP neurons are believed to show some combination of spatially selective visual, delay, or presaccadic activity during this task. However, not all neurons in LIP show these properties.

#### 4.2.4 Dimming detection task

This task was used to familiarize both animals with their familiar image set, by repeatedly presenting images from the familiar image set while maintaining the animal’s engagement.

The task begins with a fixation period, then a sequence of images is presented in the fovea. The animal is required to maintain fixation throughout the sequence. Each image is presented for 450 ms. On half the trials, there is an equal chance that 1 through 5 images will be presented. For all of these sequence lengths, the last image will dim and the animal has 450 ms to indicate that they perceived the dimming by releasing a touch bar to receive a reward. On the other half of trials, a sequence of 6 images is presented and none of them dim. To receive reward, the animal must hold the bar through the entire sequence.

### 4.3 Image sets

Both animals had a single set of familiar images that remained constant throughout the experiments. In Monkey R, this set consisted of 55 images, but 40 images were randomly selected for use in each experimental session. In Monkey N, this set consisted of 40 images and the whole set was used in each experimental session. All image sets (both novel and familiar) for each experimental session across both monkeys was randomly selected from the Corel Image Library (which includes photographs of a variety of natural scenes, including a variety of objects, buildings, and animals). The images were cropped to be 150×150 pixels. These images were not controlled for their low-level features or for their contrast, as doing so often made them appear unnatural and we believe that this would introduce a bigger confound than attempting to control for these properties. We evaluated the effect of low-level image salience on animal viewing behavior in the PLT and did not find a reliable relationship (see *The effect of low-level salience on behavior* in *Supplement*).

### 4.4 Electrophysiological recording

In Monkey R, neuronal activity was recorded using 75 μm tungsten microelectrodes (FHC) or 16-channel V-Probes (Plexon). In Monkey N, neuronal recordings were made using either 16- or 24-channel V-Probes (Plexon). All 16-channel probes had 100 μm spacing between electrode sites; 24-channel probes had 75 μm spacing. In both cases, all of the sites were arranged in a line. In Monkey R, 25 of the sessions used single wire electrodes (FHC). In the rest of the sessions in Monkey R and all of the sessions in Monkey N, we used 16- or 24-channel probes (Plexon). During recordings, one of the two test images in both tasks was placed within the spatial response field (RF) of as many neurons as possible recorded in that session. The spatial response field was determined through the standard memory-guided saccade task (see *Memory-guided saccade* in *Methods* for task details). Neurophysiological signals from both single or multi-channel recording were amplified, digitized and stored for offline spike sorting. In both monkeys, we recorded neuronal activity in the memory-guided saccade task to map LIP RF locations.

We localized LIP in each monkey according to the pattern of neuronal activity during the memory-guided saccade task, as described above. In both monkeys, we also considered neurons recorded from the same grid holes and at similar depths as previous recordings that yielded LIP-like memory-guided saccade activity to also be in LIP. In all recordings, we also identified LIP neurons based on anatomical criteria, such as the location of each electrode track relative to that expected from the MRI scans, the pattern of gray–white matter transitions encountered on each electrode penetration, and the relative depths of each neuron.

#### 4.4.1 Spike sorting

In Monkey R, 44/66 of the datasets were sorted offline by hand (Plexon) to verify the quality and stability of neuronal isolations. In the remaining 22/66 of the datasets from Monkey R and for all of the data from Monkey N, single neurons were sorted automatically using Kilosort 2 (code available [here](https://github.com/MouseLand/Kilosort2))[8]. All analyses were performed on neurons marked “good” by that software, which indicates that less than 10% of recorded spikes are likely to have arisen from a different neuron. Postprocessing of these sorted neurons was performed in Phy (code available [here](https://github.com/cortex-lab/phy/)).

#### 4.4.2 Preprocessing

For all analyses reported here, the data from each neuron was smoothed with a 50 ms wide causal boxcar filter. That is, the spike rate at time zero includes all of the spikes from –50 - 0 ms. Where noted, we also z-scored these firing rates.

### 4.5 Data analysis

#### 4.5.1 Exclusion criteria

Only sessions in which monkeys performed enough trials for each unique stimulus condition in both the sDMST and PLT tasks (*n* ≥ 30 for monkeys R and N) were used for further analysis, except in figs. 4 and 5 where neurons with *n* ≥ 20 trials were included (since there were fewer neurons with sufficient trials for the conPLT). Further, in pseudo-population analyses, only trials in which the neuron fired more than 2 spikes through the whole trial were included. The neuron totals for each pseudo-population analysis are reported in the main text. In addition, for true population decoding analyses, only recordings with more than five neurons are included.

#### 4.5.2 Saccade detection and analysis

To detect saccades, we first applied a median filter to the recorded eye traces to remove noise. Then, saccades were defined as periods within the smoothed eye trace during which the instantaneous velocity of the eye was greater than 70 ° s^−1^. Any fixations that lasted less than 10 ms were assumed to be artifactual and therefore removed so that the previous and succeeding saccades were merged into a single saccade. Velocity for the whole saccade was computed by taking the distance between the eye position at the beginning and end of the saccade, divided by the duration of the saccade.

We organize our analyses around the time of the animal’s first saccade in the response period (that is, after the fixation point disappears in both tasks, and the free viewing array appears in the PLT and the test array appears in the sDMST). Time zero in all of the plots with time on the x-axis is the time of the initiation of that saccade.

#### 4.5.3 Support vector machine (SVM) analyses

We trained support vector machine (SVM) classifiers to decode both the animal’s saccade choice (i.e., a saccade into or away from the RF of the recorded neurons) and information about the image in the RF. In all decoding analyses the number of trials in each condition were balanced. Further, when decoding information about the image in the RF, an even number of trials where the animal made a saccade into and away from the RF were included - thus, marginalizing over the animal’s motor action. These balanced sets of trials were resampled 500 times to account for the variability that is introduced by choosing only a subset of trials. These 500 resamplings were used to produce 95% confidence intervals for our SVM plots, as displayed.

All of our SVM analyses depend on the SVM implementation provided in scikit-learn[9]. We used the radial basis function kernel in all cases. Performance was not qualitatively different when using a linear kernel for all the analyses.

#### 4.5.4 Data and code availability

All code used to generate the figures and perform the analyses is written in python and available [here](https://github.com/wj2/vplt-analysis). This code relies on additional custom code available [here](https://github.com/wj2/general-neural). Finally, it also makes extensive use of the python scientific computing environment.

## 5 Supplement

### 5.1 Additional behavioral quantification in the PLT

Here, we describe an additional way of evaluating the animal’s familiarity bias as well as investigate the animal’s image-independent bias toward a particular side. As reported in the main text, on most experimental sessions both animals exhibited a significant familiarity bias in their first saccade (Figure S.1a). Another way of looking at this evaluates the amount of time that the animal spends viewing the novel relative to the familiar image in the early part of the PLT free viewing period. That is, we take the 100 - 350 ms after the onset of the free viewing period, which encapsulates most of the animal’s first and second fixation. Then, we take the amount of time their eyes were on the familiar image and subtract that by the amount of time their eyes were on the novel image. Then, we divide by the period length (250 ms). This gives us a looking time familiarity bias that ranges from −1 to 1, where −1 indicates that the animal spent all of their time viewing the familiar image and 1 indicates that they spent all of their time viewing the novel image. Both animals have significant, and quite large, familiarity biases according to this metric as well (Monkey R: 0.26 - 0.39; Monkey N: –0.52 - –0.24; Figure S.1b).

Finally, we also compute the animal’s side bias for their first saccade. That is, we ask if the animal is more likely to look to the image in one hemifield independent of any of that image’s properties. Both animal’s have often significant side biases that vary across sessions (mean side bias from different experimental sessions, Monkey R: 0.43 - 0.54, Monkey N: 0.26 - 0.4; Figure S.1c). Though in many behavioral sessions, this bias is relatively small.

Next, we ask about the interactions between these three quantities. First, we ask whether the animal’s first saccade bias and looking time bias are correlated with each other across sessions. That is, when the animal has a strong first saccade bias, do they also have a strong looking time bias? Interestingly, we do not find a reliable relationship between these two quantities (Monkey R: *r* =–0.54 - 0.06, Monkey N: *r* =–0.48 - 0.47; Figure S.1d). Next, we asked if the strength of the animal’s side bias predicted the strength of the animal’s first saccade bias. Here, we do find a strong negative relationship in both monkeys (Monkey R: *r* =–0.73 –0.3, Monkey N: *r* =–0.86 –0.36; Figure S.1e). Together, these results indicate that the animal’s side bias may influence that strength of the animal’s first saccade bias. However, the strength of the first saccade bias is not strongly related to the animal’s overall familiarity preference.

**Figure S.1:**
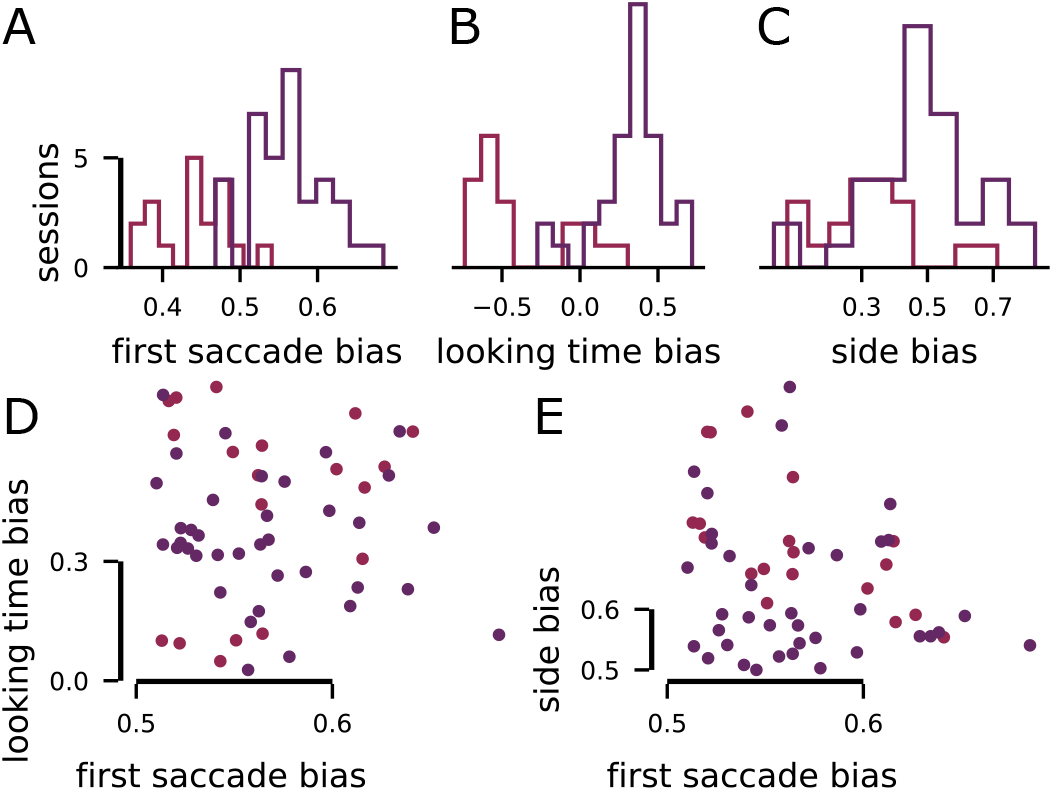
Additional quantification of the animal’s behavior on the PLT, related to Figure 1. **A** Histogram of the first saccade novelty bias across different experimental sessions. **B** Histogram of the looking time novelty bias across different experimental sessions. **C** Histogram of the side bias across different experimental sessions. Here, 1 means the animal always looked into the recorded hemifield, and 0 means the animal always looked away from the recorded hemifield. **D** Scatter plot between the first saccade bias and looking time bias. This plot indicates no significant relationship between these two quantities. **E** Scatter plot between the first saccade bias and the side bias. This plot reveals that, as the side bias becomes larger, the first saccade bias tends to decrease.

### 5.2 The effect of low-level salience on behavior

One possibility is that the animal’s behavior in the PLT is strongly entrained by differences in the low-level salience of the two presented images. Low-level salience is derived from areas within each image of high contrast or high spatial frequency relative to the surroundings. Many algorithms have been proposed to compute this low-level salience from image inputs, and are benchmarked against human free viewing data[1]. One successful yet simple algorithm is Boolean map salience (BMS)[2]. BMS leverages both local salience cues like those described above, as well as some global cues that have also been shown to be used in figure-ground segregation such as surroundedness.

We evaluated whether the animal’s behavior was significantly modulated by differences in image salience. To do this, we computed the low-level salience map of each of the images in the familiar image set, using the BMS algorithm, and tested whether the animal was more likely to look at the image with higher mean salience either with their first saccade (Figure S.2a) or for more time in the initial part of the free-viewing period (Figure S.2b) – both of which are time periods where the animal’s behavior shows significant modulation by differences in familiarity. For both ways of computing salience bias, neither animal showed a reliable bias toward the more salient images across different experimental sessions. Further, the biases that were expressed in each session were almost always smaller in magnitude than the familiarity bias and likely reflect random fluctuations in the animal’s behavior rather than a true salience bias (looking time bias, Monkey R: –0.04 - 0.02 difference in looking time; first saccade bias, Monkey R: 0.47 - 0.52 probability of looking to the more salient image).

**Figure S.2:**
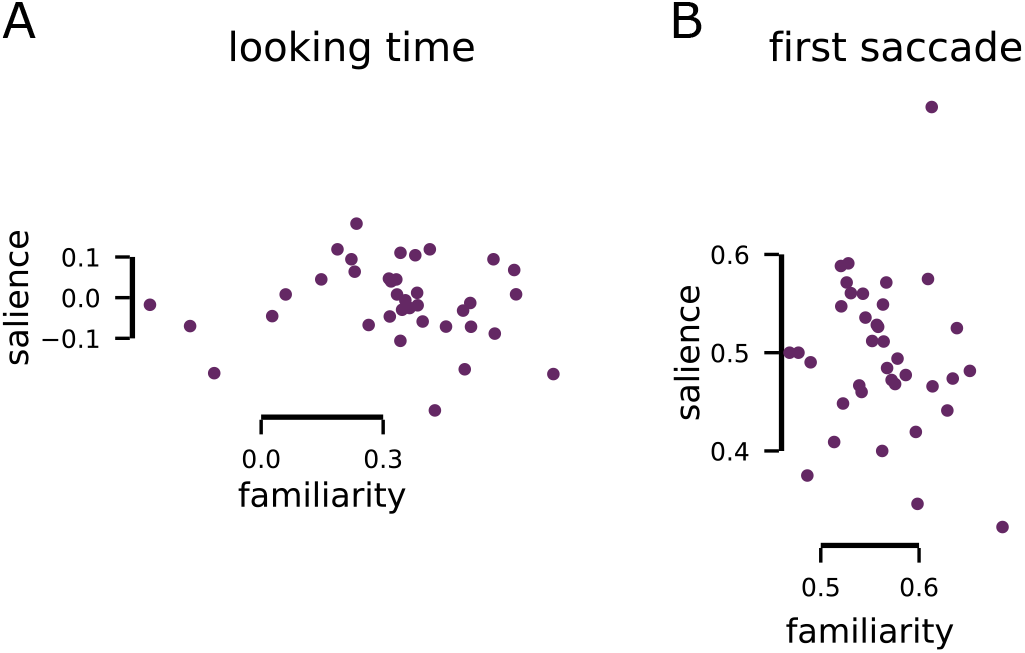
Quantification of the animal’s behavior as it relates to the low-level salience of the presented images, related to Figure 4. **A** Scatter plot showing the relationship between the strength of the biasing of the first saccade due to differences in low-level salience or differences in familiarity. There is no significant relationship. **B** Scatter plot showing the relationship between the strength of the looking time bias due to differences in low-level saliences and differences in familiarity. There is no significant relationship.

### 5.3 Undirected tasks with stimuli in close spatial proximity

In addition to the conditions of the tasks focused on in the main text, we included conditions in both the PLT and lumPLT in which the two images were placed with 45° rather than 180° angular separation. This condition allows us to ask if LIP’s representation of either the behavioral response or of image relevance changes when the two images are placed in close proximity to each other. As low-level salience and other methods of capturing bottom-up attention are often defined relative to immediately surrounding stimuli, this closer stimulus proximity in our task might reveal a more direct role for LIP in undirected behavior.

As in the main text, we train SVM classifiers to decode both the saccade choice (Figure S.3a,b) and image relevance (Figure S.3c,d) when the images are placed close and far from each other. We perform this analysis on both the standard PLT (Figure S.3a,c) and the luminance PLT (Figure S.3b,d). In all cases, the results are similar between the close and far conditions, indicating that LIP does not play a special role for undirected stimulus selection among nearby rather than far apart stimuli.

**Figure S.3:**
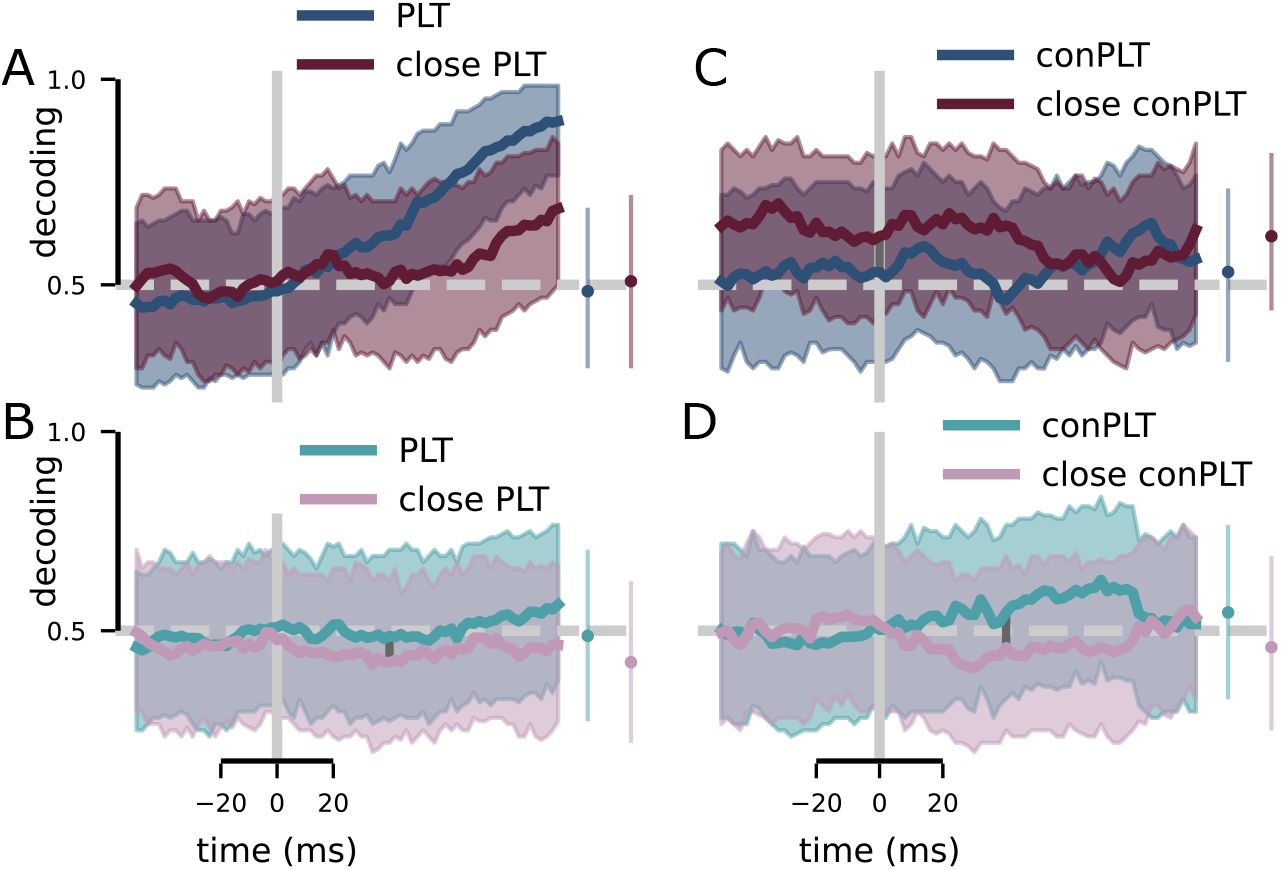
Comparison of decoding performance when stimuli are in close proximity relative to when they are in opposite hemifields, related to Figure 5. **A** Decoding performance for saccade choice in the standard PLT where stimuli were placed either in opposite hemifields or nearby. **B** Decoding performance for saccade choice in the luminance PLT where stimuli were placed either in opposite hemifields or nearby. **C** Decoding performance for familiarity in the standard PLT where stimuli were placed either in opposite hemifields or nearby. **D** Decoding performance for contrast level in the luminance PLT where stimuli were placed either in opposite hemifields or nearby.

## References

1. Desimone, R. & Duncan, J. Neural mechanisms of selective visual attention. Annual review of neuroscience 18, 193–222 (1995).

2. Ungerleider, S. K. & G, L. Mechanisms of visual attention in the human cortex. Annual review of neuroscience 23, 315–341 (2000).

3. Teller, D. Y. The forced-choice preferential looking procedure: A psychophysical technique for use with human infants. Infant Behavior and Development 2, 135–153 (1979).

4. Jutras, M. J. & Buffalo, E. A. Recognition memory signals in the macaque hippocampus. Proceedings of the National Academy of Sciences 107, 401–406 (2010).

5. Manns, J. R., Stark, C. E. & Squire, L. R. The visual paired-comparison task as a measure of declarative memory. Proceedings of the National Academy of Sciences 97, 12375–12379 (2000).

6. Pascalis, O. & Bachevalier, J. Face recognition in primates: a cross-species study. Behavioural processes 43, 87–96 (1998).

7. Bichot, N. P., Schall, J. D. & Thompson, K. G. Visual feature selectivity in frontal eye fields induced by experience in mature macaques. 1996.

8. Horowitz, T. S. & Wolfe, J. M. Visual search has no memory. Nature 394, 575–577 (1998).

9. Wardak, C., Olivier, E. & Duhamel, J.-R. Saccadic target selection deficits after lateral intraparietal area inactivation in monkeys. The Journal of neuroscience : the official journal of the Society for Neuroscience 22, 9877–84. ISSN: 1529-2401. http://www.ncbi.nlm.nih.gov/pubmed/12427844 (2002).

10. Buschman, T. J. & Miller, E. K. Top-down versus bottom-up control of attention in the prefrontal and posterior parietal cortices. Science (New York, N.Y.) 315, 1860–1862. ISSN: 0036-8075 (2007).

11. Reynolds, J. H. & Chelazzi, L. Attentional modulation of visual processing. Annu. Rev. Neurosci. 27, 611–647 (2004).

12. Ipata, A. E., Gee, A. L., Goldberg, M. E. & Bisley, J. W. Activity in the Lateral Intraparietal Area Predicts the Goal and Latency of Saccades in a Free-Viewing Visual Search Task. The Journal of Neuroscience 26, 3656–3661. ISSN: 0270-6474, 1529-2401 (2006).

13. Ipata, A. E., Gee, A. L., Bisley, J. W. & Goldberg, M. E. Neurons in the lateral intraparietal area create a priority map by the combination of disparate signals. Experimental brain research 192, 479–488 (2009).

14. Kusunoki, M., Gottlieb, J. & Goldberg, M. E. The lateral intraparietal area as a salience map: The representation of abrupt onset, stimulus motion, and task relevance. Vision Research 40, 1459–1468. ISSN: 00426989 (2000).

15. Foley, N. C., Jangraw, D. C., Peck, C. & Gottlieb, J. Novelty enhances visual salience independently of reward in the parietal lobe. J Neurosci 34, 7947–7957. ISSN: 1529-2401. http://www.ncbi.nlm.nih.gov/pubmed/24899716 (2014).

16. Bisley, J. W. & Mirpour, K. The neural instantiation of a priority map. Current opinion in psychology 29, 108–112 (2019).

17. Schiller, P., Tehovnik, E. & Larry R. Squire. Cortical Control of Eye Movements, 175–181 (2009).

18. Ibos, G., Duhamel, J.-R. & Hamed, S. B. A functional hierarchy within the parietofrontal network in stimulus selection and attention control. Journal of Neuroscience 33, 8359–8369 (2013).

19. Chen, X. et al. Parietal Cortex Regulates Visual Salience and Salience-Driven Behavior. Neuron (2020).

20. Freedman, D. J. & Assad, J. A. Experience-dependent representation of visual categories in parietal cortex. Nature 443, 85 (2006).

21. Grunewald, A., Linden, J. F. & Andersen, R. A. Responses to auditory stimuli in macaque lateral intraparietal area I. Effects of training. Journal of neurophysiology 82, 330–342 (1999).

22. Sarma, A., Masse, N. Y., Wang, X.-J. & Freedman, D. J. Task-specific versus generalized mnemonic representations in parietal and prefrontal cortices. Nature neuroscience 19, 143–149. ISSN: 1546-1726 (2016).

23. Ibos, G. & Freedman, D. J. Dynamic integration of task-relevant visual features in posterior parietal cortex. Neuron 83, 1468–1480 (2014).

24. Zhou, Y. & Freedman, D. J. Posterior parietal cortex plays a causal role in perceptual and categorical decisions. Science 365, 180–185 (2019).

25. Swaminathan, S. K. & Freedman, D. J. Preferential encoding of visual categories in parietal cortex compared with prefrontal cortex. Nature neuroscience 15, 315–320 (2012).

26. Barash, S., Bracewell, R. M., Fogassi, L., Gnadt, J. W. & Andersen, R. A. Saccade-related activity in the lateral intraparietal area. I. Temporal properties; comparison with area 7a. J Neurophysiol 66, 1095–1108. ISSN: 00223077. http://www.ncbi.nlm.nih.gov/pubmed/1753276 (1991).

27. Barash, S., Bracewell, R. M., Fogassi, L., Gnadt, J. W. & Andersen, R. a. Saccade-related activity in the lateral intraparietal area. II. Spatial properties. Journal of neurophysiology 66, 1109–1124. ISSN: 00223077 (1991).

28. Li, C.-S. R., Mazzoni, P. & Andersen, R. A. Effect of reversible inactivation of macaque lateral intraparietal area on visual and memory saccades. Journal of Neurophysiology 81, 1827–1838 (1999).

29. Ipata, A. E., Gee, A. L., Gottlieb, J., Bisley, J. W. & Goldberg, M. E. LIP responses to a popout stimulus are reduced if it is overtly ignored. Nature neuroscience 9, 1071–1076 (2006).

30. Fahy, F., Riches, I. & Brown, M. Neuronal activity related to visual recognition memory: longterm memory and the encoding of recency and familiarity information in the primate anterior and medial inferior temporal and rhinal cortex. Experimental Brain Research 96, 457–472 (1993).

31. Huang, G., Ramachandran, S., Lee, T. S. & Olson, C. R. Neural correlate of visual familiarity in macaque area V2. Journal of Neuroscience 38, 8967–8975 (2018).

32. Freedman, D. J., Riesenhuber, M., Poggio, T. & Miller, E. K. Experience-dependent sharpening of visual shape selectivity in inferior temporal cortex. Cerebral Cortex 16, 1631–1644 (2006).

33. Anderson, B., Mruczek, R. E. B., Kawasaki, K. & Sheinberg, D. Effects of familiarity on neural activity in monkey inferior temporal lobe. Cerebral Cortex 18, 2540–2552. ISSN: 10473211. arXiv: NIHMS150003 (2008).

34. Woloszyn, L. & Sheinberg, D. L. L. Effects of Long-Term Visual Experience on Responses of Distinct Classes of Single Units in Inferior Temporal Cortex. Neuron 74, 193–205. ISSN: 08966273 (2012).

35. Mountcastle, V. B., Lynch, J., Georgopoulos, A., Sakata, H. & Acuna, C. Posterior parietal association cortex of the monkey: command functions for operations within extrapersonal space. Journal of neurophysiology 38, 871–908 (1975).

36. Lynch, J., Mountcastle, V., Talbot, W. & Yin, T. Parietal lobe mechanisms for directed visual attention. Journal of Neurophysiology 40, 362–389 (1977).

37. Robinson, D. L., Goldberg, M. E. & Stanton, G. B. Parietal association cortex in the primate: sensory mechanisms and behavioral modulations. journal of Neurophysiology 41, 910–932 (1978).

38. Gnadt, J. W. & Andersen, R. A. Memory related motor planning activity in posterior parietal cortex of macaque. Experimental brain research 70, 216–220 (1988).

39. Ungerleider, L. G. & Mishkin, M. Two cortical visual systems 1982.

40. Mishkin, M., Ungerleider, L. G. & Macko, K. A. Object vision and spatial vision: two cortical pathways. Trends in Neurosciences 6, 414–417 (1983).

## References

1. Swaminathan, S. K., Masse, N. Y. & Freedman, D. J. A comparison of lateral and medial intraparietal areas during a visual categorization task. Journal of Neuroscience 33, 13157–13170 (2013).

2. Rishel, C. A., Huang, G. & Freedman, D. J. Independent category and spatial encoding in parietal cortex. Neuron 77, 969–979. http://dx.doi.org/10.1016/j.neuron.2013.01.007 (2013).

3. Asaad, W. F., Santhanam, N., McClellan, S. & Freedman, D. J. High-performance execution of psychophysical tasks with complex visual stimuli in MATLAB. Journal of neurophysiology 109, 249–260 (2013).

4. Hwang, J., Mitz, A. R. & Murray, E. A. NIMH MonkeyLogic: Behavioral control and data acquisition in MATLAB. Journal of neuroscience methods 323, 13–21 (2019).

5. Gnadt, J. W. & Andersen, R. A. Memory related motor planning activity in posterior parietal cortex of macaque. Experimental brain research 70, 216–220 (1988).

6. Roitman, J. D. & Shadlen, M. N. Response of neurons in the lateral intraparietal area during a combined visual discrimination reaction time task. Journal of neuroscience 22, 9475–9489 (2002).

7. Hikosaka, O. & Wurtz, R. H. Visual and oculomotor functions of monkey substantia nigra pars reticulata. III. Memory-contingent visual and saccade responses. Journal of neurophysiology 49, 1268–1284 (1983).

8. Pachitariu, M., Steinmetz, N., Kadir, S., Carandini, M. & Harris, K. Fast and accurate spike sorting of high-channel count probes with KiloSort, 1–9 (2016).

9. Pedregosa, F. et al. Scikit-learn: Machine Learning in Python. Journal of Machine Learning Research 12, 2825–2830 (2011).

## References

1. Borji, A., Cheng, M.-M., Jiang, H. & Li, J. Salient object detection: A benchmark. IEEE transactions on image processing 24, 5706–5722 (2015).

2. Zhang, J. & Sclaroff, S. Saliency detection: A boolean map approach in Proceedings of the IEEE international conference on computer vision (2013), 153–160.

